# Preferential targeting of lateral entorhinal inputs onto newly integrated granule cells

**DOI:** 10.1101/153767

**Authors:** Nicholas I. Woods, Christopher E. Vaaga, Christina Chatzi, Jaimie D. Adelson, Matthew F. Collie, Julia V. Perederiy, Kenneth R. Tovar, Gary L. Westbrook

## Abstract

Mature dentate granule cells in the hippocampus receive input from the entorhinal cortex via the perforant path in precisely arranged lamina, with medial entorhinal axons innervating the middle molecular layer and lateral entorhinal cortex axons innervating the outer molecular layer. Although vastly outnumbered by mature granule cells, adult-generated newborn granule cells play a unique role in hippocampal function, which has largely been attributed to their enhanced excitability and plasticity (Schmidt-Hieber et al., 2004; Ge et al., 2007). Inputs from the medial and lateral entorhinal cortex carry different informational content, thus the distribution of inputs onto newly integrated granules will affect their function in the circuit. Therefore we examined the functional innervation and synaptic maturation of newly-generated dentate granule cells using retroviral labeling in combination with selective optogenetic activation of medial or lateral entorhinal inputs. Our results indicate that lateral entorhinal inputs provide nearly all the functional innervation of newly integrated granule cells. Despite preferential functional targeting, the dendritic spine density of immature granule cells was not increased in the outer molecular layer compared to the middle molecular layer. However, chronic blockade of neurotransmitter release in medial entorhinal axons with tetanus toxin disrupted normal synapse development from both medial and lateral entorhinal inputs. Our results support a role for preferential lateral perforant path input onto newly generated neurons in mediating pattern separation, but also indicates that medial perforant path input is necessary for normal synaptic development.

**Significance Statement:** The formation of episodic memories involves the integration of contextual and spatial information. Newly integrated neurons in the dentate gyrus of the hippocampus play a critical role in this process, despite constituting only a minor fraction of total granule cells. Here we demonstrate that these neurons preferentially receive information thought to convey the context of an experience - a unique role that each newly integrated granule cell serves for about a month before reaching maturity.

## Introduction

As the entry point to the trisynaptic hippocampal circuit, the dentate gyrus has several interesting features including a ‘sparse’ network design (Boss et al., 1985; Rolls et al., 1998), laminated inputs carrying distinct informational content (Witter, 2007; Knierim et al., 2014), and participation of mature granule cells alongside the continuous integration of newly-generated neurons (Overstreet-Wadiche and Westbrook, 2006; Ming and Song, 2011). Hippocampal granule cells receive highly laminar inputs from entorhinal cortex within the molecular layer of the dentate gyrus. Input from medial entorhinal cortex, conveying spatial cues, is restricted to the middle molecular layer (Ferbinteanu et al., 1999; Hafting et al., 2005; Hargreaves et al., 2005; Yasuda and Mayford, 2006; Witter, 2007; Van Cauter et al., 2013), whereas input from lateral entorhinal cortex conveying contextual information is restricted to the outer molecular layer (Hargreaves et al., 2005; Hunsaker et al., 2007; Witter, 2007; Deshmukh and Knierim, 2011; Yoganarasimha et al., 2011; Tsao et al., 2013). Despite being substantially outnumbered by mature granule cells, newly integrated granule cells are thought to uniquely contribute to pattern separation (Clelland et al., 2009; Sahay et al., 2011; Nakashiba et al., 2012; Tronel et al., 2012) - that is, the ability to distinguish between subtly different contexts - one of the primary functions of the dentate gyrus (Deng et al., 2010; Aimone et al., 2011). This unique function must occur during a narrow time window between initial integration into the perforant path circuit (~3 weeks post-mitosis) and complete maturation of synapses (>8 weeks) (van Praag et al., 2002; Ambrogini et al., 2004; Overstreet et al., 2004; Overstreet-Wadiche et al., 2006; Zhao et al., 2006; Brunner et al., 2014).

The search for unique functions of newly integrated neurons has largely focused on intrinsic properties, such as enhanced excitability and plasticity (Schmidt-Hieber et al., 2004; Abrous et al., 2005; Ge et al., 2007; Marín-Burgin et al., 2012; Dieni et al., 2013). However, newborn neurons also undergo rapid changes in connectivity, which differs from synapse remodeling in early development (Goodman and Shatz, 1993; Katz and Shatz, 1996; Walsh and Lichtman, 2003), as newborn neurons integrate into an already established circuit (Toni et al., 2007, 2008), and compete for synaptic innervation with pre-existing axons of the perforant path. Rabies-based circuit mapping suggests that inputs to newly integrated neurons may differ from mature granule cells (Vivar et al., 2012).

Here, we directly assayed synaptic integration of newborn granule cells over the course of excitatory synapse development using retroviral labeling of newborn neurons and laminar-specific optogenetic stimulation of entorhinal inputs. Our results indicate that newly integrated granule cells preferentially receive functional synaptic input from lateral entorhinal cortex, whereas mature granule cells receive balanced input from medial and lateral entorhinal cortex. Although medial perforant path input was weak in newly integrated granule cells, chronic silencing of this pathway using tetanus toxin impaired the functional and morphological development of lateral perforant path inputs.

## Methods

### Animals

We used male and female C57/Bl6 mice. Animal procedures were carried out in accordance with the Oregon Health and Science University Institutional Animal Care and Use Committee, Biosafety Committee protocols, and NIH guidelines for the safe handling of animals.

### Viral Constructs

To selectively transfect and visualize adult-born hippocampal granule cells, we used a replication-deficient Moloney Murine Leukemia Virus-based retroviral vector that requires cell mitosis for transduction (Luikart et al., 2011). The retrovirus contained an internal ubiquitin 6 promoter that drives expression of GFP (viral titer 10^5^) as described previously (Luikart et al., 2011). To express the light-activated ion channel channelrhodopsin-2 (ChR2) selectively in entorhinal cortex axons projecting to the middle or outer molecular layer, we stereotaxically injected an AAV9-CAG-ChR2-eGFP viral construct (UNC Viral Core) into the medial or lateral entorhinal cortex. To selectively silence axons, we injected a custom AAV-CAG-TeNT-mCherry virus (viral titer 10^13^) made by cloning a 2kb fragment encoding the light chain of tetanus toxin fused with mCherry into an AAV backbone using InFusion cloning. AAV vectors were serotyped with AAV9 coat proteins and packaged at the University of North Carolina Vector Core.

### Stereotaxic Injections

Stereotaxic viral injections into 6-8 week old male and female C57/Bl6 mice were carried out using a Model 1900 Stereotaxic Alignment System (Kopf). Mice were anesthetized with 2% isoflurane, and a small incision was created over the skull following application of artificial tears to the eyes and antibiotic/iodine around the incision site. A Model 1911 Stereotaxic Drill was used to create burr holes over the injection site. pRubi-GFP retrovirus to label mitotically active granule cells was injected into the dentate gyrus (coordinates: in mm from bregma): anteroposterior: −1.9, lateromedial: +/- 1.1; dorsoventral: −2.5, −2.3. One microliter of non-diluted virus was injected at 250 nl/min with a 10 μl Hamilton syringe fitted with a 30-gauge needle using a Quintessential stereotaxic injector (Stoetling). After each injection, the needle was left in place for 1 minute to allow for diffusion of the virus and prevent backflow. Injections of AAV9-CAG-CHR2-GFP into the lateral entorhinal cortex were made at anteroposterior: −3.4, lateromedial: +/- 4.0; dorsoventral: −2.4. Injections of AAV9-CAG-ChR2-GFP or AAV9-CAG-TeNT-mCherry into the medial entorhinal cortex were made at anteroposterior: −4.5; dorsoventral: +/−3.0; dorsoventral: −3.2. Following injection, mice received topical lidocaine and drinking water with cherry-flavored Tylenol, and monitored every 24 hours over 3 days.

### Immunohistochemistry and Spine Counts

Mice were transcardially perfused with ice-cold 4% sucrose in 0.1 M PBS followed by fixative containing 4% sucrose and 4% paraformaldehyde (PFA) in PBS. Brains were removed, and postfixed (4% PFA, 1x PBS) overnight at 4°C. The dorsal hippocampus was sectioned (coronal, 100 μm) using a Leica VT 1000s vibratome. Sections were permeabilized with 0.4% Triton X-100 in PBS (PBS-T), blocked with filtered 10% horse serum in PBS-T and incubated in primary antibody overnight in 1.5% horse serum in PBS-T. Primary antibodies included: 1:300 Alexa-Fluor 488-conjugated rabbit anti-GFP (catalog no. A21311, Invitrogen), 1:500 anti-VGlut2 (catalog no. 135 404, Synaptic Systems), and 1:500 anti-glial associated fibrillary protein (GFAP; catalog no. Z-0334, DAKO). Sections were then rinsed with PBS-T and incubated for 2-3 hours at room temperature in PBS-T with secondary antibodies: 1:300 Alexa Fluor 488-conjugated anti-GFP (A21311, Invitrogen), 1:200 goat anti-guinea pig Alexa Fluor 568 (A11075, Invitrogen), 1:200 goat anti-rabbit Alexa Fluor 488). The mCherry-TeNT signal was visualized using native fluorescence. The tissue was counterstained with DAPI using Fluoromount G with DAPI (SouthernBiotech).

Images were acquired using either a Zeiss LSM 770 or LSM 780 laser-scanning confocal microscope on a motorized AxioObserver Z1 inverted scope (Carl Zeiss MicroImaging). Dendrites for spinal analysis were imaged using a 63x objective (1.4 NA, oil, 2x zoom). For each imaged cell, dendritic segments in the middle molecular layer and the outer molecular layer were imaged. The middle and outer molecular layer were distinguished based on VGluT2 immunofluorescence pattern, which begins at the border between the inner molecular layer and middle molecular layer. Middle molecular layer dendritic segments were therefore imaged at the beginning of the VGluT2 staining, whereas outer molecular layer dendritic segments were imaged at the distal tip of the molecular layer. Spine density analysis was performed blinded to experimental condition. The Cell Counter plugin in FIJI (NIH) was used to count and categorize spines (Harris et al., 1992), and Simple Neurite Tracer (FIJI) was used to measure dendritic segment length. Spine density and spine type between the middle and outer molecular layer was compared across conditions (control and TeNT overexpression) and developmental time points (3-12 weeks post-viral injection). Spine morphology was visually assessed: dendritic spines containing a spine head (ca. 2x shaft diameter) were considered as mushroom spines and all other spines were considered filopodia-like.

### Electrophysiology

Electrophysiological recordings were made 21 days after viral injection to allow for construct expression. Acute coronal brain slices were prepared as described previously (Perederiy et al., 2013). Briefly, animals were anesthetized with an intraperitoneal injection of 2% 2,2,2-tribromoethanol (0.7-0.8 mL), and transcardially perfused with an ice-cold, oxygenated modified ACSF which contained (in mM): 110 choline-Cl, 7 MgCl2, 2.5 KCl, 1.25 NaH2PO4, 0.5 CaCl, 1.3 Na-ascorbate, and 25 NaHCO3. Hippocampi were resected and cut at 300 μm in the transverse axis on a Leica 1200s vibratome. Slices were allowed to incubate for 1 hr in 37°C normal ACSF, which contained (in mM): 125 NaCl, 2.5 KCl, 2.0 CaCl, 1.0 MgCl2, 1.25 NaH2PO4, 25 NaHCO3, and 25 glucose.

We used glass pipettes (2-3 MΩ) filled with normal ACSF for extracellular field recording, which were placed into the lamina of interest. Presynaptic fibers were stimulated using a bipolar electrode (3-7v, 0.5v steps) or optogenetic stimulation (1ms pulses of 470 nm blue light). Whole-cell voltage clamp recordings were made using glass pipettes (5-8 MΩ). Mature granule cells were selected based on input resistance less than 750 MΩ (495±37 MΩ) and soma position in the outer ⅓ of the granule cell layer (Ambrogini et al., 2004; Overstreet-Wadiche and Westbrook, 2006). The whole-cell recording solution contained (in mM): 100 gluconic acid, 0.2 EGTA, 5 HEPES, 2 Mg-ATP, 0.3 Li-GTP (pH: 7.2, 295 mOsm; adjusted with 50% CsOH such that final concentration of Cs-gluconate is 100-120 mM). The liquid junction potential (- 7 mV) was not corrected. Input resistance of the cell was continually monitored with a 10 mV hyperpolarizing step, and cells with input resistance exceeding 25 MΩ at any point were excluded from analysis. Data was acquired at 10 KHz and Bessel filtered at 4 KHz on a Multiclamp 700B (Axon Instruments, Sunnyvale CA) and recorded using AxographX acquisition software (www.axograph.com).

Optogenetic stimulation was provided by an 470 nm LED (ThorLabs, Newton, NJ). Light pulses (1 ms) were provided through the microscope objective, which was centered over the appropriate lamina. Optical stimulation was provided over a range of intensities until a maximal response was elicited, which was then used for the remainder of the experiment. Peak EPSC amplitudes were measured using a built-in routine in AxographX. Miniature EPSCs were recorded in the presence of SR95531 (10 μM) and TTX (1 μM) to isolate miniature excitatory events. Quantal events were detected using a sliding window template consisting of a single exponential (-10 pA, 1 ms rise time, 6 ms decay time constant). Individual events were then manually inspected. mEPSC analysis was performed with the experimenter blinded to condition.

### Cell Culture

Mouse hippocampal neurons were cultured on glial micro-islands as described previously (Tovar et al., 2009). Briefly, neonatal (postnatal day 0-1) male mice were decapitated, and the hippocampi were dissected. Micro-islands were generated by plating at 125,000 cells/35 mm dish. After 7 days *in vitro*, cultures were treated with 200 μM glutamate for 30 min to kill any neurons. Neurons were then plated on the remaining glial feeder layer at 25,000 cells/35 mm dish and maintained in a tissue culture incubator (37°C, 5% CO_2_) until use. The culture medium consisted of minimum essential media with 2 mM glutaMAX (Invitrogen), 5% heat-inactivated fetal calf serum (Lonza), and 1 ml/l MITO+ Serum Extender (BD Biosciences). The culture medium was supplemented with glucose to a final concentration of 21 mM. Cultured neurons were transduced at 1 day *in vitro* by replacing 50% of the culture medium with virus-containing medium (1 μL of virus in 500 μL medium). After 24 hours, the virus-containing medium was removed and replaced with fresh complete medium.

Whole cell voltage clamp recordings were made from cultured neurons 3-16 days *in vitro*. The extracellular recording solution consisted of (in mM): 158 NaCl, 2.4 KCl, 1.3 CaCl_2_, 1 MgCl_2_, 10 HEPES and 10 D-glucose (pH 7.4; 320 mosmol). Glass pipettes (2-6 MΩ) were filled with a solution which contained (in mM): 140 K-gluconate, 4 CaCl_2_, 8 Na Cl, 2 MgCl_2_, 10 EGTA, 10 HEPES, 4 Na_2_ATP and 0.2 Na_2_GTP (pH: 7.4, 319 mosmol). Autaptic EPSCs were elicited from neurons in isolation on a glial micro-island with a brief voltage command (+30 mV, 0.5 ms) to elicit an unclamped action potential. Recordings were made in the presence of 10 μM SR95531 and 10 μM (R)-CPP to block GABAA and NMD_A_ receptors, respectively. Data was acquired using an Axopatch 1C amplifier and AxographX (www.axograph.com) acquisition software. In all recordings, the series resistance was <10 MΩ and was continuously monitored with a −10 mV step. Data was low-pass Bessel filtered at 4 kHz and sampled at 10 kHz.

### Electron Microscopy

Ultrastructural analysis of control and TeNT expressing medial perforant path axons was done 21-days post viral injection, as in Perederiy et al., 2013. Two control (pRubi-expressing) and two pRubi and TeNT-expressing animals were transcardially perfused with PBS followed by a 3.75% acrolein and 2% paraformaldehyde fixative. The brains were then extracted and stored in 2% paraformaldehyde for at least 1 hour prior to sectioning at 40 μm in the coronal plane using a Leica VT 1000s vibratome (Leica Microsystems). Sections including the dorsal hippocampus were incubated in 1% sodium borohydride for 30 minutes to reduce nonspecific binding, followed by incubation in 10% Triton-X for 45 minutes to increase antibody penetration. Next, sections were blocked with 0.5% bovine serum albumin for 1 hour followed by primary antibody incubation directed against pRubi-GFP (Rabbit α-GFP; 1:500, Millipore Cat #: AB3080) or TeNT-mCherry (Mouse α-mCherry; 1-500, Living Colors Cat #: 632543) overnight at 4°C. Following primary antibody incubation, the tissue was thoroughly washed with 0.4% Triton-X. To visualize GFP, tissue was incubated in biotinylated goat α-rabbit secondary antibody (1:200; Vector Laboratories, Cat # BA-1000) for 2 hours at room temperature followed by avidin-binding complex (Vector Laboratories, Burlingame CA) for 30 minutes then reacted with DABH_2_O_2_ solution for 5.5 minutes. To visualize mCherry, the tissue was incubated in goat α-mouse gold-conjugated IgG (1:50; Aurion, Cat #: 800.422) for 2 hours at room temperature. Tissue was then washed with citrate buffer and silver enhanced for 6.5 minutes.

Following DAB and/or immunogold reactions, the tissue was fixed in 1% osmium tetroxide for 15 minutes in 0.1 M phosphate buffer. Tissue was then washed and dehydrated through an ethanol series before being incubated in propylene oxide (10 minutes) and propylene oxide:EMBed (1:1 solution) overnight. Finally, the tissue was embedded in Aclar resin and placed in an oven at 60°C for 24 hours. 700 nm coronal sections were made using a Leica EM UC6 vibratome (Leica Microsystems). Some sections were mounted on glass slides and stained with toluidine blue in 0.5 % sodium tetraborate to assist in region selection. Tissue from the supra-pyramidal blade of the dentate gyrus was sectioned at 70 nm using an ultramicrotome (Leica Microsystems). Sections were placed on 200 square mesh copper/rhodium grids and counterstained with 5% uranyl acetate and Reynold’s lead citrate. The middle molecular layer was imaged at 11,000x on an FEI Technai G^2^ 12 BioTWIN microscope at 80 kV. At least 10 representative images were selected in both control and TeNT expressing conditions.

### Experimental Design and Statistical Analysis

Male and female mice were used for all experiments except developmental spine analysis, in which only males were used to provide more consistent results across animals. All data are reported as mean±SEM unless otherwise noted. Statistical analysis was performed in Prism6 (GraphPad Software, La Jolla, CA). Data were assumed to be normally distributed, in accordance with previous datasets in this circuit. Spine density data were analyzed using a two-way repeated measures ANOVA (repeated measures: days post-mitosis, lamina). Pooled data were analyzed using an ANOVA with Holm-Sidak post-hoc analysis. Electrophysiology data were analyzed using two-tailed unpaired Student’s t-tests. Linear regressions were performed using an Extra-sum of squares F-test. Sample sizes were chosen to detect an effect size of 20%, based on previous experiments, with a power of 0.8. Statistical significance was defined as α<0.05, and was adjusted for post-hoc comparisons (Holm-Sidak), as appropriate.

## Results

### Adult-born granule cells receive preferential input from outer molecular layer axons

To examine the perforant path input onto newborn granule cells, we used electrical and optogenetic laminar-specific stimulation (Figure 1 A). Targeting of the medial or lateral perforant path was possible using a bipolar electrode as demonstrated by changes in the polarity of the field EPSP (Figure 1 B, Andersen et al., 1966). However, optogenetic labeling allows more precise pathway-specific stimulation. Thus we injected AAV9-CAG-ChR2-eGFP into either medial or lateral entorhinal cortex, which provided very precise labeling of either the medial or lateral perforant path fibers in the molecular layer (Figure 1C).

**Figure 1:**
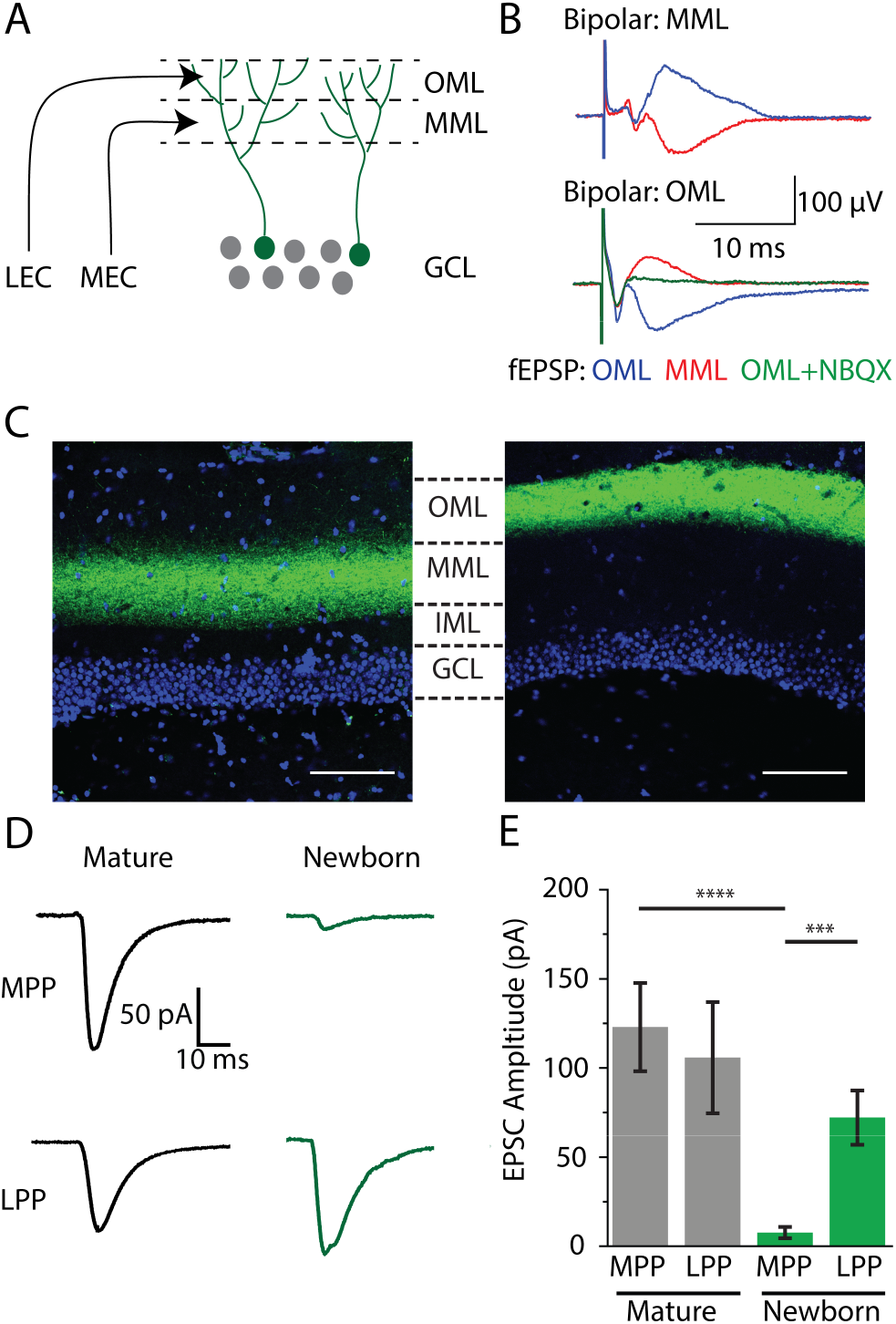
Laminar specific electrical and optical stimulation. (A) Circuit schematic demonstrating laminar specific input from medial entorhinal cortex (MEC) and lateral entorhinal cortex (LEC). Within the molecular layer of the dentate gyrus, MEC axons reside in the middle molecular layer (MML) whereas lateral entorhinal cortex axons reside in the outer molecular layer (OML). (B) Demonstration of laminar specific input using field EPSP recordings. When the field recording electrode and bipolar electrode are within the same layer, a current sink is observed as a negative voltage deflection. A current source can be observed when the field electrode and bipolar electrode are in adjacent layers. (C) Laminar specific expression of ChR2 following viral injection into the medial entorhinal cortex (left) or lateral entorhinal cortex (right). Scale bar: 100 μm. (D) Comparison of lamina-specific optogenetic stimulation in mature (black) and newborn cells (green). (E) Comparison of the strength of the maximal light evoked EPSCs from each lamina in mature and newborn (p21) granule cells.

As expected, in mature cells, there was no difference in the maximal amplitude of light-evoked EPSCs from the medial (MPP) or lateral perforant path (LPP) (MPP: 123.0±24.7 pA, n=19 cells; LPP: 105.9±31.3 pA, n=16 cells; Student’s unpaired t-test: t(33): 0.46, p=0.67 ; Figure 1 D, E), indicating that mature cells receive robust and balanced excitatory input from both layers. Surprisingly, in newly integrated neurons (retrovirally labeled cells, 21 days post-mitosis), the strength of lateral perforant path inputs was nearly 10-fold larger than inputs from the medial perforant path (MPP: 7.8±3.1 pA, n=14 cells; LPP: 72.2±15.2 pA, n=18 cells; Student’s unpaired t-test: t(30): 3.68, p=0.0009:; Figure 1 D, E). The reduced strength of medial perforant path inputs in newly integrated granule cells was not a result of NMDA-only or ‘silent’ synapses, as there was no difference in the AMPA/NMDA ratio between lamina (MPP: 5.2±1.3, n=13 cells; LPP: 3.66±0.58, n=13 cells; Student’s unpaired t-test: t(24): 1.04; p=0.31). These results indicate that newly integrated neurons receive preferential, but not exclusive, functional input from the lateral perforant path.

### Synapse formation in newborn granule cells does not involve competitive elimination

Given the differences in synaptic strength between lateral and medial perforant path axons in newly integrated granule cells, we examined whether exuberant synapse formation occurs in the lateral perforant path, followed by competitive elimination and synaptic rebalancing, as seen during early development (Goodman and Shatz, 1993; Katz and Shatz, 1996; Walsh and Lichtman, 2003). We labeled mitotically active neurons in 6 week-old male mice with a retroviral pRUBI-GFP vector (Luikart et al. 2012), then examined spine density in 3 to 12 week-old cells (at 1 week intervals) in the middle molecular layer (MML) and outer molecular layer (OML). Despite the functional difference in synapse strength in newly integrated granule cells, there was no difference in spine density across laminae at any time point (two-way repeated measure ANOVA: F(1,2): 0.195, p=0.7). In grouped data from both laminae, the spine density increased from 3 to 4 weeks post-injection (3 wk: 0.98±0.07 spines/μm, n=36 dendritic segments from 3 animals; 4 wk: 1.52±0.03 spines/μm, n=36 dendritic segments from 3 animals; Holm-Sidak post-hoc comparison: t(47): 4.5, p<0.001; Figure 2 A, B), but then remained unchanged between 4 and 12 weeks post-injection (Holm-Sidak post-hoc comparison: p>0.05). The spine density at 12 weeks is within the range of reported values for mature granule cells (Parent et al., 2016).

**Figure 2:**
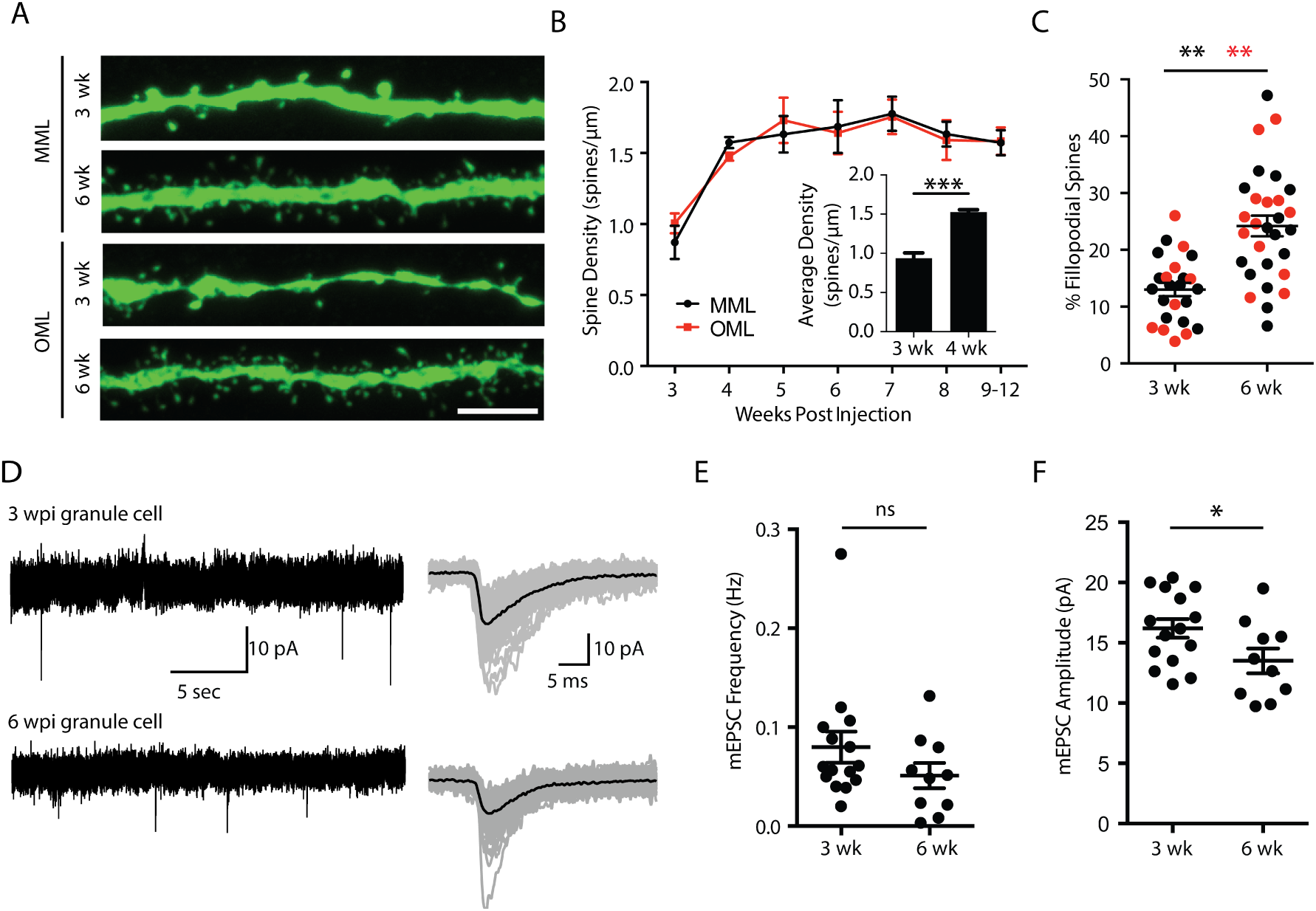
Synaptic maturation and development. (A) Spine density measurements in MML (top) and OML (bottom) at 3 and 6 week old dentate granule cells. Scale bar: 5 μm. (B) Developmental increase in spine density across cell development in middle molecular layer (black) and outer molecular layer (red). *Inset*: Average spine density across layers increases between 3 and 4 weeks post-mitosis, then remains constant between 4 and 12 weeks. (C) Developmental increase in the percentage of spines with filopodial morphology in both MML (black) and OML (red). (D) Miniature EPSC recordings in the presence of 10 μM SR95531 and 1 μM TTX to isolate excitatory events. mEPSCs were recorded at 3 and 6 weeks post mitosis. (E) There was no difference in the miniature EPSC frequency at 3 and 6 weeks post mitosis. (F) There was a significant decrease in miniature EPSC amplitude, suggesting weaker synaptic inputs at 6 weeks post-mitosis.

The morphology of dendritic spines also changed during this time period, as there was an increase in the proportion of spines with filopodial morphology in the middle molecular layer (3 wk: 13.3±1.3% of total spines, n=14 cells; 6 wk: 23.21±2.6% of total spines, n=16 cells; Student’s unpaired t-test: t(28): 3.29, p=0.0027; Figure 2 A, C) and outer molecular layer (3 wk: 12.53±2.3% of total spines, n=10 cells; 6 wk: 25.42±2.6% of total spines, n=13 cells; Student’s unpaired t-test: t(21): 3.54, p=0.002; Figure 2 A, C). In contrast to spine density, the mEPSC frequency remained unchanged between 3 and 6 weeks post mitosis (3 wk: 0.08±0.02 Hz, n=15 cells, 6 wk: 0.05±0.01 Hz, n=10 cells; Student’s unpaired t-test: t(23): 1.32, p=0.20; Figure 2 D, E). At 6 weeks post mitosis, there was a small decrease in the mESPC amplitude (3 wk: 16.2±0.8 pA, n=15 cells; 6 wk: 13.5±1.0 pA, n= 10 cells, Student’s unpaired t-test: t(23): 2.1, p=0.045, Figure 2 D, F), perhaps consistent with the observed increase in filopodial spines (Holtmaat and Svoboda, 2009). This data suggests that during this period of functional synaptic rebalancing, synapse formation by newly integrating granule cells does not involve over-abundant synapse formation followed by synaptic pruning.

### Chronic silencing with TeNT

To chronically silence axonal input in a laminar-specific manner, we virally expressed tetanus toxin light chain (TeNT), which cleaves the SNARE complex protein synaptobrevin-2, thereby preventing neurotransmitter release (Schiavo et al., 1992). We chose to use tetanus toxin because it has the advantage of completely silencing axons, with the disadvantage that effects are irreversible. As the degree of silencing may influence effects on excitatory synapse formation (Bagley and Westbrook, 2012), we carefully validated our tetanus toxin vector *in vitro* and *in vivo*. First, we examined AMPA receptor currents in neurons in autaptic cultures (Tovar et al., 2009). In control cells an unclamped action potential at the soma elicited a large amplitude, NBQX-sensitive EPSC without failure (n=7 cells; Figure 3 A, B). However, expression of TeNT by viral transfection completely abolished EPSCs (EPSC success rate: 0.22±0.2%, n=10 cells, Student’s unpaired t-test: t(14): 392.9 p<0.0001; Figure 3 A, B).

**Figure 3:**
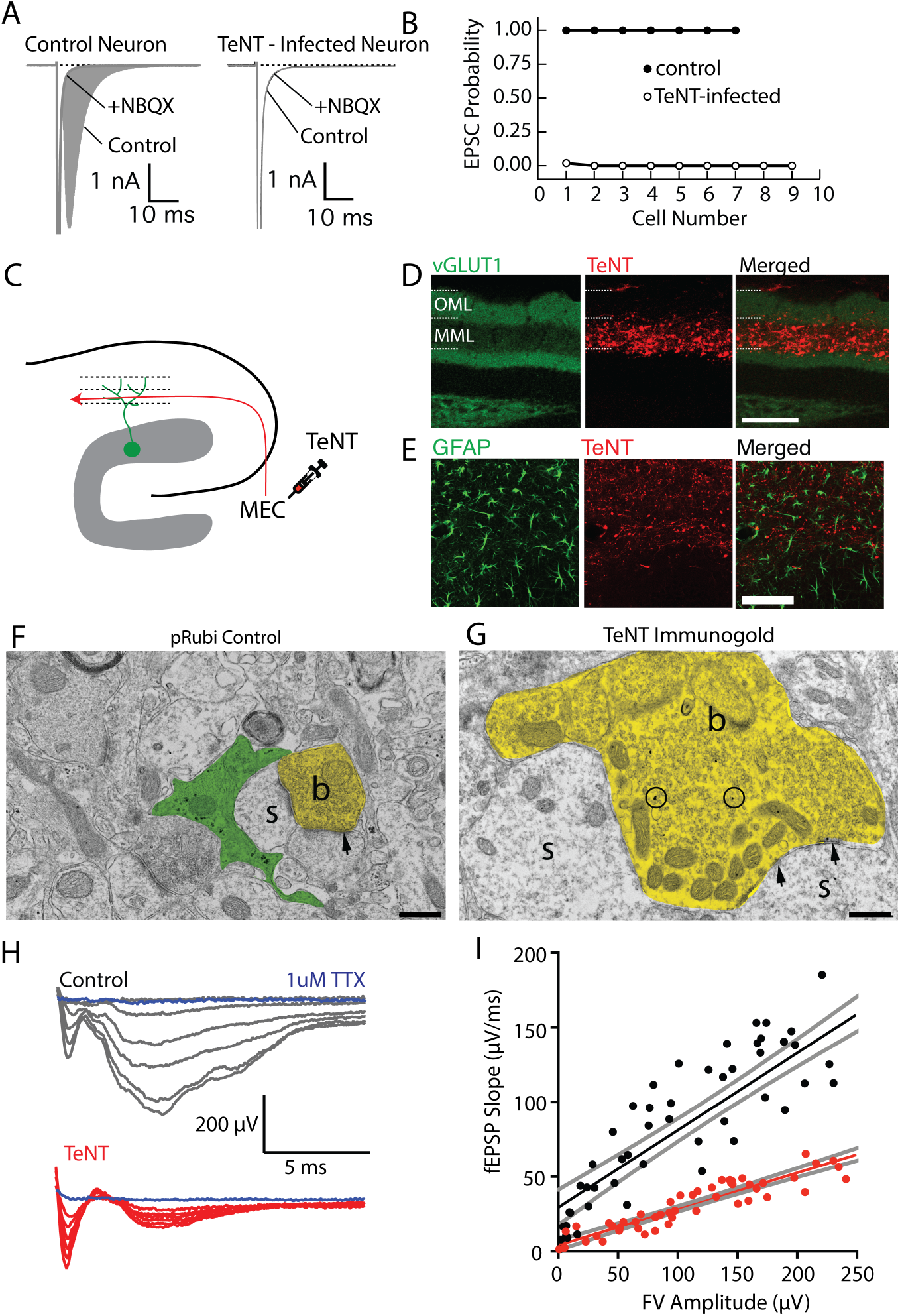
Silencing synaptic input with Tetanus toxin expression. (A) Comparison of synaptic responses in autaptically cultured neurons in control (left) and following tetanus toxin infection (TeNT). (B) Expression of TeNT completely abolishes synaptic responses in autaptically cultured neurons. (C) Schematic of TeNT viral injection into the medial entorhinal cortex, which will functionally silence axons in the middle molecular layer of the dentate gyrus. (D) Expression of TeNT in the middle molecular layer dramatically reduces the intensity of VGluT1 expression in the middle molecular layer, indicating a disruption of presynaptic function.Scale bar: 100 μm. (E) TeNT expression did not elicit astrogliosis. (F, G) Electron micrographs from control (F) and TeNT overexpressing (G) axons. TeNT expression results in axonal swelling and vesicle accumulation (b=axonal bouton, s=dendritic spine, arrowheads=synapse). Scale bar: 500 nm. (H) Field EPSP recordings from the middle molecular layer while electrically stimulating the medial perforant path fibers. (I) fEPSP responses were significantly attenuated when TeNT was expressed in the MML, without changing fiber volley amplitudes.

To selectively silence medial perforant path inputs *in vivo*, we injected an AAV9-TeNT-mCherry virus into the medial entorhinal cortex (Figure 3 C), resulting in robust laminar-specific expression of TeNT in medial perforant path axons (Figure 3 C, D). TeNT expression was accompanied by a lamina-specific decrease in VGluT1 immunostaining, a marker of functional presynaptic terminals (Figure 3 D). Importantly, TeNT expression followed up to 6 weeks *in vivo* did not elicit an inflammatory response (Figure 3 E), unlike the glial scarring that occurs following axotomy of medial perforant path axons with subsequent degeneration of terminal axons (Perederiy et al., 2013). Immunogold electron micrographs of TeNT-expressing axon terminals had pronounced swelling as well as increased accumulation of small clear vesicles in presynaptic boutons, consistent with complete block of vesicular release (Figure 3 F, G). TeNT-expressing axon terminals were directly apposed to dendritic spines, further suggesting that TeNT-expression did not elicit axon degeneration.

To examine the efficiency of silencing of the middle molecular layer, we recorded fEPSP responses to laminar-specific medial perforant path electrical stimulation in control and TeNT-expressing slices. Consistent with our *in vitro* results, TeNT markedly reduced the maximal fEPSP slope (control: 0.67±0.07 μV/ms, n=6 slices; TeNT: 0.26±0.02 μV/ms, n=5 slices; Student’s unpaired t-test: t(9): 5.4, p=0.0004; Figure 3 H), as well as the input-output relationship between fiber volley amplitude and fEPSP slope (control: 0.52±0.003; TeNT: 0.24±0.01; Extra Sum of Squares F-test: F(1,105): 30.4, p<0.0001; Figure 3 I). Importantly, there was no difference in the fiber volley amplitude with maximal stimulation (control: −231.7±36.5 μV, n=5 slices; TeNT: −197±29.8 μV, n=5 slices; Student’s unpaired t-test: t(8): 0.74, p=0.48), indicating that equal numbers of axons were stimulated in both conditions. The small residual fEPSP response observed in slices likely indicates that a few medial entorhinal cortex neurons had not been infected with TeNT.

### Silencing the middle molecular layer impairs synapse formation with perforant path inputs

Although the medial perforant path provides only weak input onto newly integrated granule cells, dendritic spines are present, and thus these inputs could have an activity-dependent effect on circuit formation. We used laminar-specific silencing with TeNT to address this issue. ChR2-eGFP was expressed in lateral perforant path axons in conjunction with TeNT-mCherry expression in the medial perforant path (MPP:TeNT) (Figure 4 A). Interestingly, expression of TeNT in the medial perforant path reduced the amplitude of lateral perforant path responses in newly integrated granule cells (control: 72.2±15.2 pA, n=18 cells; MPP:TeNT:8.7±3.7 pA, n=8 cells; Student’s unpaired t-test: t(24): 2.7, p=0.011; Figure 4 B, C), but not in mature granule cells (control: 105.9±31.3 pA, n=16 cells; MPP:TeNT: 125.7±23.24 pA, n=23 cells; Student’s unpaired t-test: t(37): 0.52, p=0.6; Figure 4 B, C). There was also an increase in the paired pulse ratio of lateral perforant path axons targeting newly integrated granule cells (control PPR: 0.9±0.08, n= 18 cells, MPP:TeNT PPR: 1.8±0.3, n=8 cells; Student’s unpaired t-test: t(24): 4.3, p=0.0003; Figure 4 B, D). This change was not observed in mature granule cells (control PPR: 1.1±0.07, n= 16 cells; MPP:TeNT PPR: 1.4±0.2, n=23 cells, Student’s unpaired t-test: t(37): 1.38, p=0.17; Figure 4 B, D).

**Figure 4:**
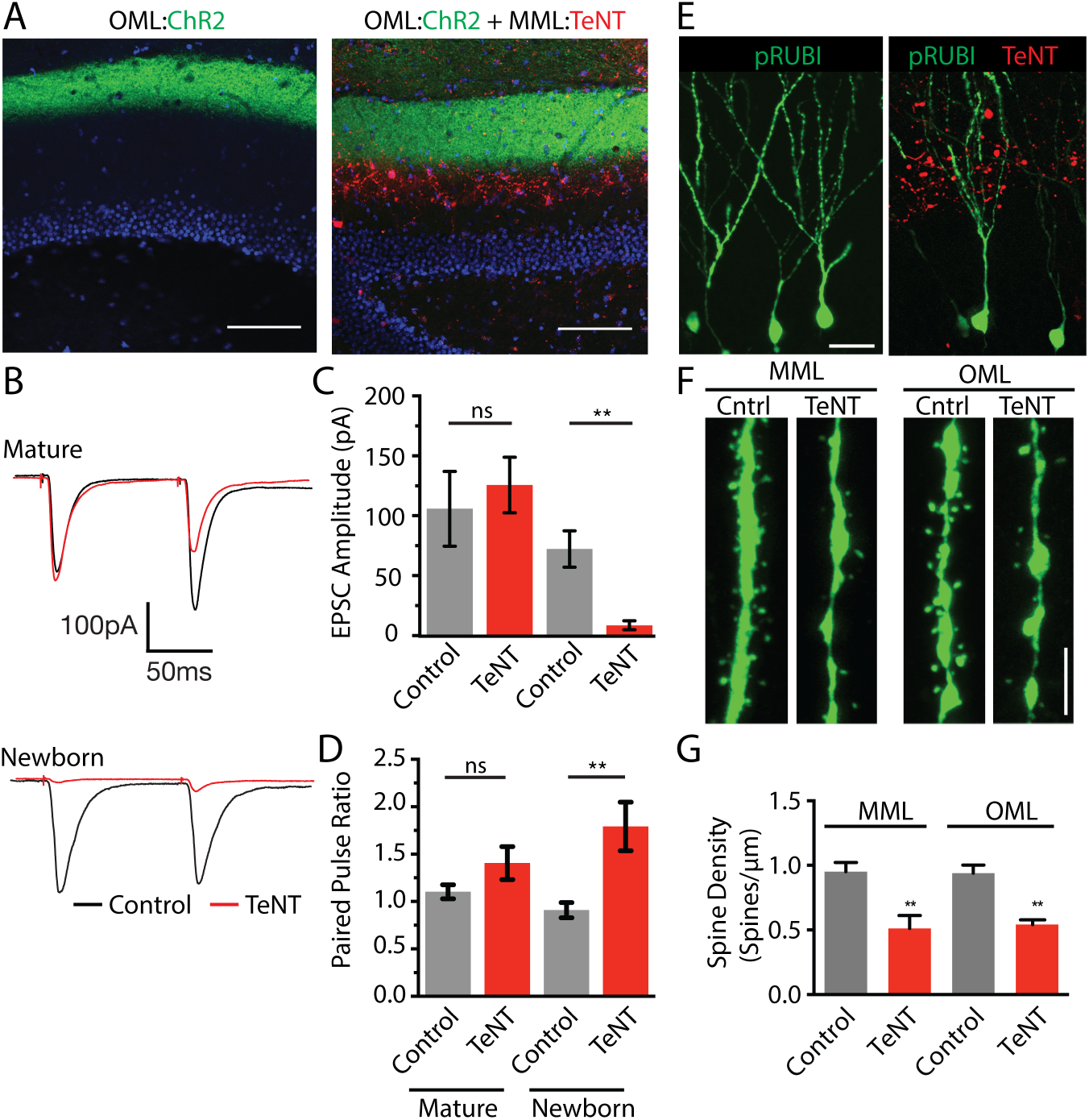
Silencing the middle molecular layer impairs normal synaptic innervation in the outer molecular layer. (A) Expression of ChR2 in the outer molecular layer, with TeNT expression in the middle molecular layer (MML:TeNT). Scale bar: 100 μm. (B) Comparison of synaptic responses in mature neurons (top) and newborn neurons (bottom) in control and MML:TeNT conditions. (C) TeNT expression in the MML significantly reduces the amplitude of OML-evoked responses selectively in newborn cells. (D) MML:TeNT expression increases the paired pulse ratio of OML axons, selectively in newborn cells. (E) Retroviral labeling of newborn dentate granule cells in control (left) and with MML:TeNT (right). Scale bar: 35 μm. (F, G) Spine density is significantly reduced in both MML and OML following MML:TeNT expression. Scale bar in (F): 2.5 μm.

Silencing of the medial perforant path also reduced the spine density of newly integrated granule cells in both the middle molecular layer (control: 0.95±0.04 spines/μm, n=24 dendritic segments from 4 animals; MPP:TeNT: 0.51±0.05 spines/μm, n=24 dendritic segments from 4 animals; Student’s unpaired t-test: t(6): 7.12, p=0.0004; Figure 4 F, G) and outer molecular layer (control: 0.93±0.03 spines/μm, n=24 dendritic segments from 4 animals; MPP:TeNT: 0.54±0.02 spines/μm, n=24 dendritic segments from 4 animals; Student’s unpaired t-test: t(6): 11.01, p<0.0001; Figure 4 F, G). Together these results suggest that although the axons of the medial perforant path make little contribution to the synaptic activation of adult-born granule cells, intact synaptic release from medial perforant path axons is required for proper functional synaptic integration of adult-born granule cells.

## Discussion

There is general consensus that newborn neurons, once they integrate into the dentate gyrus network, have a unique role in memory formation (Aimone et al., 2006; Saxe et al., 2006; Kee et al., 2007). Newborn neurons go through a relatively stereotyped maturation post-mitosis, including early GABAergic depolarization without excitatory perforant input for several weeks (Ge et al., 2006) as their dendrites extend through the molecular layer (Zhao et al., 2006). Prior results have suggested a contribution of enhanced excitability and synaptic plasticity as a reason that such a minor population can have a major impact on circuit function (Schmidt-Hieber et al., 2004; Aimone et al., 2011). However, monosynaptic labeling studies using modified rabies virus (Vivar et al., 2012) suggest that newborn neurons may have distinct connectivity as well, with extrinsic excitatory input onto newborn neurons originating primarily from the lateral entorhinal cortex. Our results indicate that newly integrated neurons receive preferential functional input from the lateral entorhinal cortex, which likely contributes to their role in pattern separation. Yet our data also show that preferential targeting must occur during a well-defined time window followed by functional synaptic reorganization that results in balanced input from medial and lateral entorhinal cortex.

### The preferential input onto newly generated granule cells

Episodic memory requires both spatial and non-spatial information, which are differentially encoded in medial and lateral entorhinal cortex, respectively (Ferbinteanu et al., 1999; Hafting et al., 2005; Hargreaves et al., 2005; Yasuda and Mayford, 2006; Hunsaker et al., 2007; Deshmukh and Knierim, 2011; Yoganarasimha et al., 2011; Tsao et al., 2013; Van Cauter et al., 2013). Thus the strict laminar organization in the molecular layer provides a framework in which distinct populations of granule cells or different regions of the dendritic tree, proximal vs distal, may differentially affect circuit function (Magee, 2000; Dieni et al., 2013, 2016). In this setting, the combination of the observed preferential functional targeting of lateral entorhinal cortex inputs onto newly integrated granule cells complements their well-documented enhanced plasticity (Schmidt-Hieber et al., 2004; Abrous et al., 2005; Ge et al., 2007) in mediating distinct aspects of memory formation (Clelland et al., 2009; Sahay et al., 2011; Nakashiba et al., 2012; Tronel et al., 2012). For example, this may fit with the role of newly integrated neurons as novelty detectors for incoming contextual information, the essence of pattern separation (Deng et al., 2010; Aimone et al., 2011). The preferential input from lateral entorhinal cortex indicates that information processing in newly integrated neurons is functionally distinct from mature neurons. Although the specific information carried by the lateral and medial perforant path is likely to be more complex than a simple segregation of spatial and contextual input (Knierim et al., 2014), the strict anatomical lamination of the molecular layer suggests that the two inputs remain segregated along the dendritic tree of granule cells.

Despite the weak input from the medial perforant path, in our experiments there was no difference in spine density on newly integrated granule cells between middle and outer molecular layers. At the synaptic level, the preferential functional input could not be attributed to an increase in ‘silent synapses’ (Isaac et al., 1995; Carroll and Malenka, 2000; Ziv and Garner, 2001; Feldman, 2009), as there was no difference in the AMPA/NMDA ratio between medial and lateral perforant path inputs. Perhaps surprisingly, the presence of the same density of spines within the middle molecular layer indicates spine morphology is dissociated from functional synaptic strength in newly generated granule cells. Although presynaptic axon terminals generally co-localize with postsynaptic spines, spine formation can be temporally distinct from functional synapse formation (Yuste and Bonhoeffer, 2004). In fact, following lesions of the perforant path, newly integrated granule cells continue to form dendritic spines despite the loss of presynaptic axon terminals (Perederiy et al., 2013; see also Sando et al., 2017). Furthermore, excitatory synapses may initially form on dendritic shafts as opposed to spines (Crain et al., 1973; Miller and Peters, 1981; Mates and Lund, 1983; Yuste and Bonhoeffer, 2004; Fortin et al., 2014), which may explain why the mEPSC frequency did not increase between 3 and 6 weeks post-mitosis in our experiments.

### The role of the medial perforant path

Our results indicate medial perforant path inputs are necessary for normal synapse formation of all perforant path inputs on newly integrated granule cells. Adult-born granule cells in the hippocampus share many properties with immature neurons during development (Schmidt-Hieber et al., 2004; Abrous et al., 2005; Ge et al., 2007), but are unique in that they must integrate into a pre-existing circuit (Ge et al., 2007; Toni et al., 2007; Adlaf et al., 2017). The preferential functional targeting by the lateral entorhinal cortex is somewhat surprising from a developmental perspective given that as newborn cells mature, their dendrites first pass through the middle molecular layer, which is occupied by axons innervating mature granule cells. Inputs from the medial entorhinal cortex were weak in newly integrated neurons, perhaps explaining why it was not detected in rabies tracing studies of 21 days post-mitosis granule cells (Vivar et al., 2012). However, chronic silencing of medial perforant path inputs with tetanus toxin nearly eliminated the strong input from the lateral entorhinal cortex without affecting the lamination of incoming axons. The effect of silencing was selective for inputs onto newly generated neurons and occurred even though not every axon in the medial perforant path expressed tetanus, as estimated from the residual field EPSP. Silencing the medial perforant path was accompanied by a reduction in spine density in both the middle and outer molecular layer, although the reduction in synaptic strength was greater than the reduction in spine density. Interestingly, this pattern contrasts with homeostatic plasticity observed in some circuits (Davis, 2013). Given the weak nature of this input, it may be that factors other than net neural activity contribute to the developmental role of the medial perforant path inputs (i.e. neurotrophic factors; (Huang and Reichardt, 2001; Cohen-Cory et al., 2010).

### Comparison to synapse formation during early development

The existence of activity- and competition-dependent synapse remodeling is well mapped in the immature brain as neural circuits first develop (Goodman and Shatz, 1993; Katz and Shatz, 1996; Walsh and Lichtman, 2003). In many developing brain circuits, neurons initially form an overabundance of weak synapses which are later pruned in a competition-dependent manner, resulting in the retention of strong synaptic inputs (Bear, 1995; Knudsen, 2004; Majewska and Sur, 2006; Bhatt et al., 2009; Feldman, 2009). Such activity-dependent synaptic competition is critical in the formation of mature, functional circuits (LeVay et al., 1980; Walsh and Lichtman, 2003; Datwani et al., 2009). These processes of synapse pruning and redistribution occur during critical periods of development, when incoming patterns of activity strongly influence circuit remodeling (Malenka and Bear, 2004; Holtmaat and Svoboda, 2009; Caroni et al., 2014). However, in the adult brain, such circuit plasticity is more limited (Tagawa et al., 2005; Sato and Stryker, 2008), suggesting that the basic pattern of initial synapse formation and subsequent remodeling/refinement in adult-born cells may be distinct. Indeed we did not see a period of synaptic overabundance as newly integrated cells reached maturity. Rather the period of synaptic ‘competition’ for newly integrating neurons reflects a rebalancing of functional inputs across the molecular layer. Remodeling in the adult environment is relevant not only to neurogenic niches, but also to repair after neural injury or cell transplantation approaches (Lindvall and Kokaia, 2006; Lepousez et al., 2015).

## Acknowledgements

This work was supported by R01 NS080979 and the Ellison Medical Foundation (GLW) and P30 NS061800. We thank Stefanie Kaech Petrie for help with imaging. Thanks to Eric Schnell for guidance with electrophysiology. We thank Sue Aicher for assistance with electron microscopy.

## References

Abrous DN, Koehl M, Le Moal M (2005) Adult neurogenesis: from precursors to network and physiology. Physiol Rev 85:523–569.

Adlaf EW, Vaden RJ, Niver AJ, Manuel AF, Onyilo VC, Araujo MT, Dieni CV, Vo HT, King GD, Wadiche JI, Overstreet-Wadiche L (2017) Adult-born neurons modify excitatory synaptic transmission to existing neurons. Elife 6 Available at: http://dx.doi.org/10.7554/eLife.19886.

Aimone JB, Deng W, Gage FH (2011) Resolving new memories: a critical look at the dentate gyrus, adult neurogenesis, and pattern separation. Neuron 70:589–596.

Aimone JB, Wiles J, Gage FH (2006) Potential role for adult neurogenesis in the encoding of time in new memories. Nat Neurosci 9:723–727.

Ambrogini P, Lattanzi D, Ciuffoli S, Agostini D, Bertini L, Stocchi V, Santi S, Cuppini R (2004) Morpho-functional characterization of neuronal cells at different stages of maturation in granule cell layer of adult rat dentate gyrus. Brain Res 1017:21–31.

Bagley EE, Westbrook GL (2012) Short-term field stimulation mimics synaptic maturation of hippocampal synapses. J Physiol 590:1641–1654.

Bear MF (1995) Mechanism for a sliding synaptic modification threshold. Neuron 15:1–4.

Bhatt DH, Zhang S, Gan W-B (2009) Dendritic spine dynamics. Annu Rev Physiol 71:261–282.

Boss BD, Peterson GM, Cowan WM (1985) On the number of neurons in the dentate gyrus of the rat. Brain Res 338:144–150.

Brunner J, Neubrandt M, Van-Weert S, Andrási T, Kleine Borgmann FB, Jessberger S, Szabadics J (2014) Adult-born granule cells mature through two functionally distinct states. Elife 3:e03104.

Caroni P, Chowdhury A, Lahr M (2014) Synapse rearrangements upon learning: from divergent-sparse connectivity to dedicated sub-circuits. Trends Neurosci 37:604–614.

Carroll RC, Malenka RC (2000) Delivering the goods to synapses. Nat Neurosci 3:1064–1066.

Clelland CD, Choi M, Romberg C, Clemenson GD Jr, Fragniere A, Tyers P, Jessberger S, Saksida LM, Barker RA, Gage FH, Bussey TJ (2009) A functional role for adult hippocampal neurogenesis in spatial pattern separation. Science 325:210–213.

Cohen-Cory S, Kidane AH, Shirkey NJ, Marshak S (2010) Brain-derived neurotrophic factor and the development of structural neuronal connectivity. Dev Neurobiol 70:271–288.

Crain B, Cotman C, Taylor D, Lynch G (1973) A quantitative electron microscopic study of synaptogenesis in the dentate gyrus of the rat. Brain Res 63:195–204.

Datwani A, McConnell MJ, Kanold PO, Micheva KD, Busse B, Shamloo M, Smith SJ, Shatz CJ (2009) Classical MHCI molecules regulate retinogeniculate refinement and limit ocular dominance plasticity. Neuron 64:463–470.

Davis GW (2013) Homeostatic signaling and the stabilization of neural function. Neuron 80:718–728.

Deng W, Aimone JB, Gage FH (2010) New neurons and new memories: how does adult hippocampal neurogenesis affect learning and memory? Nat Rev Neurosci 11:339–350.

Deshmukh SS, Knierim JJ (2011) Representation of non-spatial and spatial information in the lateral entorhinal cortex. Front Behav Neurosci 5:69.

Dieni CV, Nietz AK, Panichi R, Wadiche JI, Overstreet-Wadiche L (2013) Distinct determinants of sparse activation during granule cell maturation. J Neurosci 33:19131–19142.

Dieni CV, Panichi R, Aimone JB, Kuo CT, Wadiche JI, Overstreet-Wadiche L (2016) Low excitatory innervation balances high intrinsic excitability of immature dentate neurons. Nat Commun 7:11313.

Feldman DE (2009) Synaptic mechanisms for plasticity in neocortex. Annu Rev Neurosci 32:33–55.

Ferbinteanu J, Holsinger RM, McDonald RJ (1999) Lesions of the medial or lateral perforant path have different effects on hippocampal contributions to place learning and on fear conditioning to context. Behav Brain Res 101:65–84.

Fortin DA, Tillo SE, Yang G, Rah J-C, Melander JB, Bai S, Soler-Cedeño O, Qin M, Zemelman BV, Guo C, Mao T, Zhong H (2014) Live imaging of endogenous PSD-95 using ENABLED: a conditional strategy to fluorescently label endogenous proteins. J Neurosci 34:16698–16712.

Ge S, Goh ELK, Sailor KA, Kitabatake Y, Ming G-L, Song H (2006) GABA regulates synaptic integration of newly generated neurons in the adult brain. Nature 439:589–593.

Ge S, Yang C-H, Hsu K-S, Ming G-L, Song H (2007) A critical period for enhanced synaptic plasticity in newly generated neurons of the adult brain. Neuron 54:559–566.

Goodman CS, Shatz CJ (1993) Developmental mechanisms that generate precise patterns of neuronal connectivity. Cell 72 Suppl:77–98.

Hafting T, Fyhn M, Molden S, Moser M-B, Moser EI (2005) Microstructure of a spatial map in the entorhinal cortex. Nature 436:801–806.

Hargreaves EL, Rao G, Lee I, Knierim JJ (2005) Major dissociation between medial and lateral entorhinal input to dorsal hippocampus. Science 308:1792–1794.

Harris KM, Jensen FE, Tsao B (1992) Three-dimensional structure of dendritic spines and synapses in rat hippocampus (CA1) at postnatal day 15 and adult ages: implications for the maturation of synaptic physiology and long-term potentiation [published erratum appears in J Neurosci 1992 Aug; 12 (8): following table of contents]. Journal of Neuroscience 12:2685–2705.

Holtmaat A, Svoboda K (2009) Experience-dependent structural synaptic plasticity in the mammalian brain. Nat Rev Neurosci 10:647–658.

Huang EJ, Reichardt LF (2001) Neurotrophins: roles in neuronal development and function. Annu Rev Neurosci 24:677–736.

Hunsaker MR, Mooy GG, Swift JS, Kesner RP (2007) Dissociations of the medial and lateral perforant path projections into dorsal DG, CA3, and CA1 for spatial and nonspatial (visual object) information processing. Behav Neurosci 121:742–750.

Isaac JT, Nicoll RA, Malenka RC (1995) Evidence for silent synapses: implications for the expression of LTP. Neuron 15:427–434.

Katz LC, Shatz CJ (1996) Synaptic activity and the construction of cortical circuits. Science 274:1133–1138.

Kee N, Teixeira CM, Wang AH, Frankland PW (2007) Preferential incorporation of adult-generated granule cells into spatial memory networks in the dentate gyrus. Nat Neurosci 10:355–362.

Knierim JJ, Neunuebel JP, Deshmukh SS (2014) Functional correlates of the lateral and medial entorhinal cortex: objects, path integration and local-global reference frames. Philos Trans R Soc Lond B Biol Sci 369:20130369.

Knudsen EI (2004) Sensitive periods in the development of the brain and behavior. J Cogn Neurosci 16:1412–1425.

Lepousez G, Nissant A, Lledo P-M (2015) Adult neurogenesis and the future of the rejuvenating brain circuits. Neuron 86:387–401.

LeVay S, Wiesel TN, Hubel DH (1980) The development of ocular dominance columns in normal and visually deprived monkeys. J Comp Neurol 191:1–51.

Lindvall O, Kokaia Z (2006) Stem cells for the treatment of neurological disorders. Nature 441:1094–1096.

Luikart BW, Schnell E, Washburn EK, Bensen AL, Tovar KR, Westbrook GL (2011) Pten knockdown in vivo increases excitatory drive onto dentate granule cells. J Neurosci 31:4345–4354.

Magee JC (2000) Dendritic integration of excitatory synaptic input. Nat Rev Neurosci 1:181–190.

Majewska AK, Sur M (2006) Plasticity and specificity of cortical processing networks. Trends Neurosci 29:323–329.

Malenka RC, Bear MF (2004) LTP and LTD: an embarrassment of riches. Neuron 44:5–21.

Marín-Burgin A, Mongiat LA, Pardi MB, Schinder AF (2012) Unique processing during a period of high excitation/inhibition balance in adult-born neurons. Science 335:1238–1242.

Mates SL, Lund JS (1983) Spine formation and maturation of type 1 synapses on spiny stellate neurons in primate visual cortex. J Comp Neurol 221:91–97.

Miller M, Peters A (1981) Maturation of rat visual cortex. II. A combined Golgi-electron microscope study of pyramidal neurons. J Comp Neurol 203:555–573.

Ming G-L, Song H (2011) Adult neurogenesis in the mammalian brain: significant answers and significant questions. Neuron 70:687–702.

Nakashiba T, Cushman JD, Pelkey KA, Renaudineau S, Buhl DL, McHugh TJ, Rodriguez Barrera V, Chittajallu R, Iwamoto KS, McBain CJ, Fanselow MS, Tonegawa S (2012) Young dentate granule cells mediate pattern separation, whereas old granule cells facilitate pattern completion. Cell 149:188–201.

Overstreet LS, Hentges ST, Bumaschny VF, de Souza FSJ, Smart JL, Santangelo AM, Low MJ, Westbrook GL, Rubinstein M (2004) A transgenic marker for newly born granule cells in dentate gyrus. J Neurosci 24:3251–3259.

Overstreet-Wadiche LS, Bensen AL, Westbrook GL (2006) Delayed development of adult-generated granule cells in dentate gyrus. J Neurosci 26:2326–2334.

Overstreet-Wadiche LS, Westbrook GL (2006) Functional maturation of adult-generated granule cells. Hippocampus 16:208–215.

Parent AS, Pinson A, Woods N, Chatzi C, Vaaga CE, Bensen A, Gérard A, Thome JP, Bourguignon JP, Westbrook GL (2016) Early exposure to Aroclor 1254 in vivo disrupts the functional synaptic development of newborn hippocampal granule cells. Eur J Neurosci 44:3001–3010.

Perederiy JV, Luikart BW, Washburn EK, Schnell E, Westbrook GL (2013) Neural injury alters proliferation and integration of adult-generated neurons in the dentate gyrus. J Neurosci 33:4754–4767.

Rolls ET, Treves A, Rolls ET (1998) Neural networks and brain function. Available at: http://www.oxcns.org/papers/Rolls%20Treves%201998%20Capacity%20of%20pattern%20ssociation%20networks.pdf.

Sahay A, Scobie KN, Hill AS, O’Carroll CM, Kheirbek MA, Burghardt NS, Fenton AA, Dranovsky A, Hen R (2011) Increasing adult hippocampal neurogenesis is sufficient to improve pattern separation. Nature 472:466–470.

Sando R, Bushong E, Zhu Y, Huang M, Considine C, Phan S, Ju S, Uytiepo M, Ellisman M, Maximov A (2017) Assembly of Excitatory Synapses in the Absence of Glutamatergic Neurotransmission. Neuron 94:312–321.e3.

Sato M, Stryker MP (2008) Distinctive features of adult ocular dominance plasticity. J Neurosci 28:10278–10286.

Saxe MD, Battaglia F, Wang J-W, Malleret G, David DJ, Monckton JE, Garcia ADR, Sofroniew MV, Kandel ER, Santarelli L, Others (2006) Ablation of hippocampal neurogenesis impairs contextual fear conditioning and synaptic plasticity in the dentate gyrus. Proceedings of the National Academy of Sciences 103:17501–17506.

Schiavo G, Benfenati F, Poulain B, Rossetto O, Polverino de Laureto P, DasGupta BR, Montecucco C (1992) Tetanus and botulinum-B neurotoxins block neurotransmitter release by proteolytic cleavage of synaptobrevin. Nature 359:832–835.

Schmidt-Hieber C, Jonas P, Bischofberger J (2004) Enhanced synaptic plasticity in newly generated granule cells of the adult hippocampus. Nature 429:184–187.

Tagawa Y, Kanold PO, Majdan M, Shatz CJ (2005) Multiple periods of functional ocular dominance plasticity in mouse visual cortex. Nat Neurosci 8:380–388.

Toni N, Laplagne DA, Zhao C, Lombardi G, Ribak CE, Gage FH, Schinder AF (2008) Neurons born in the adult dentate gyrus form functional synapses with target cells. Nat Neurosci 11:901–907.

Toni N, Teng EM, Bushong EA, Aimone JB, Zhao C, Consiglio A, van Praag H, Martone ME, Ellisman MH, Gage FH (2007) Synapse formation on neurons born in the adult hippocampus. Nat Neurosci 10:727–734.

Tovar KR, Maher BJ, Westbrook GL (2009) Direct actions of carbenoxolone on synaptic transmission and neuronal membrane properties. J Neurophysiol 102:974–978.

Tronel S, Belnoue L, Grosjean N, Revest J-M, Piazza P-V, Koehl M, Abrous DN (2012) Adult-born neurons are necessary for extended contextual discrimination. Hippocampus 22:292– 298.

Tsao A, Moser M-B, Moser EI (2013) Traces of experience in the lateral entorhinal cortex. Curr Biol 23:399–405.

Van Cauter T, Camon J, Alvernhe A, Elduayen C, Sargolini F, Save E (2013) Distinct roles of medial and lateral entorhinal cortex in spatial cognition. Cereb Cortex 23:451–459.

van Praag H, Schinder AF, Christie BR, Toni N, Palmer TD, Gage FH (2002) Functional neurogenesis in the adult hippocampus. Nature 415:1030–1034.

Vivar C, Potter MC, Choi J, Lee J-Y, Stringer TP, Callaway EM, Gage FH, Suh H, van Praag H (2012) Monosynaptic inputs to new neurons in the dentate gyrus. Nat Commun 3:1107.

Walsh MK, Lichtman JW (2003) In vivo time-lapse imaging of synaptic takeover associated with naturally occurring synapse elimination. Neuron 37:67–73.

Witter MP (2007) The perforant path: projections from the entorhinal cortex to the dentate gyrus. Prog Brain Res 163:43–61.

Yasuda M, Mayford MR (2006) CaMKII activation in the entorhinal cortex disrupts previously encoded spatial memory. Neuron 50:309–318.

Yoganarasimha D, Rao G, Knierim JJ (2011) Lateral entorhinal neurons are not spatially selective in cue-rich environments. Hippocampus 21:1363–1374.

Yuste R, Bonhoeffer T (2004) Genesis of dendritic spines: insights from ultrastructural and imaging studies. Nat Rev Neurosci 5:24–34.

Zhao C, Teng EM, Summers RG Jr, Ming G-L, Gage FH (2006) Distinct morphological stages of dentate granule neuron maturation in the adult mouse hippocampus. J Neurosci 26:3–11.

Ziv NE, Garner CC (2001) Principles of glutamatergic synapse formation: seeing the forest for the trees. Curr Opin Neurobiol 11:536–543.

